# A Dynamic Threshold-Based Method for Robust and Accurate Blink Detection in Eye-Tracking Data

**DOI:** 10.1101/2025.04.21.649751

**Authors:** Mohammad Ahsan Khodami

## Abstract

Blink detection is a critical component of eye-tracking research, particularly in pupillometry, where data loss due to blinks can obscure meaningful insights. Existing methods often rely on fixed thresholds or device-specific noise profiles, which may lead to inaccuracies in detecting blink onsets and offsets, especially in heterogeneous datasets. This study introduces a novel blink detection model that dynamically adapts to varying pupil size distributions, ensuring robustness across different experimental conditions. The proposed method integrates dynamic thresholding, which adjusts based on the mean pupil size of valid samples, Gaussian smoothing, which reduces noise while preserving signal integrity, and adaptive boundary refinement, which refines blink onsets and offsets based on a trends in the smoothed data. Unlike traditional approaches that merge closely spaced blinks, this model treats each blink as an independent event, preserving temporal resolution, which is essential for cognitive and perceptual studies. The model is computationally efficient and adaptable to a wide range of sampling rates, from low-frequency (e.g., 250 Hz) to high-frequency (e.g., 2000 Hz) data, ensuring consistent blink detection across different eye-tracking setups, making it suitable for both real-time and offline eye-tracking applications. Experimental evaluations demonstrate its ability to accurately detect blinks across diverse datasets. By offering a more reliable and generalizable solution, this model advances blink detection methodologies and enhances the quality of eye-tracking data analysis across research domains.

## 1 Introduction

Eye tracking technology has emerged as a fundamental methodological tool in various scientific disciplines, offering unique insights into multiple dimensions of human cognition, attention, and decision-making processes [1–8]. By recording the movements of the eyes and changes in pupil size, eye tracking technology allows researchers to objectively quantify visual engagement and explore underlying neural and psychological mechanisms [9–11]. This technology has found broad applications, including understanding reading behaviors and attentional patterns [9], decoding emotional states [12, 13], informing clinical practices [14], and examining decision-making strategies [1]. The reliability of eye-tracking data has advanced with the development of sophisticated devices, making it a foundation of modern research. However, eye-tracking data’s intrinsic complexity and variability demands accurate preprocessing to ensure that analyses yield correct and meaningful results, particularly when addressing data disruptions caused by eye blinks.

Blinks, natural physiological events characterized by rapid eyelid closure and reopening (for an extensive review, see [15]), present a notable challenge to the integrity and continuity of eye-tracking data [16, 17]. In most eye-tracking devices, blinks are identified when the measured pupil size suddenly drops to zero [18–20]. This occurs during natural blinking, as the eyelids mask the pupil [21], or due to missing data when the system does not detect the pupil accurately. Such failures may result from incorrect device calibration, participants momentarily moving out of the tracking range, or other technical limitations, leading to lost pupil signals. These interruptions, which are registered as zero pupil size, frequently cause the loss of concurrent measures (e.g., gaze position), thus creating gaps that require careful handling. Figure 1 illustrates a natural blink, and Figure 2 highlights an extreme instance of a long blink, which is not physiological but arises from factors such as the participant momentarily looking away, drifting out of the range of the device, or recording malfunctions.

**Fig. 1.**
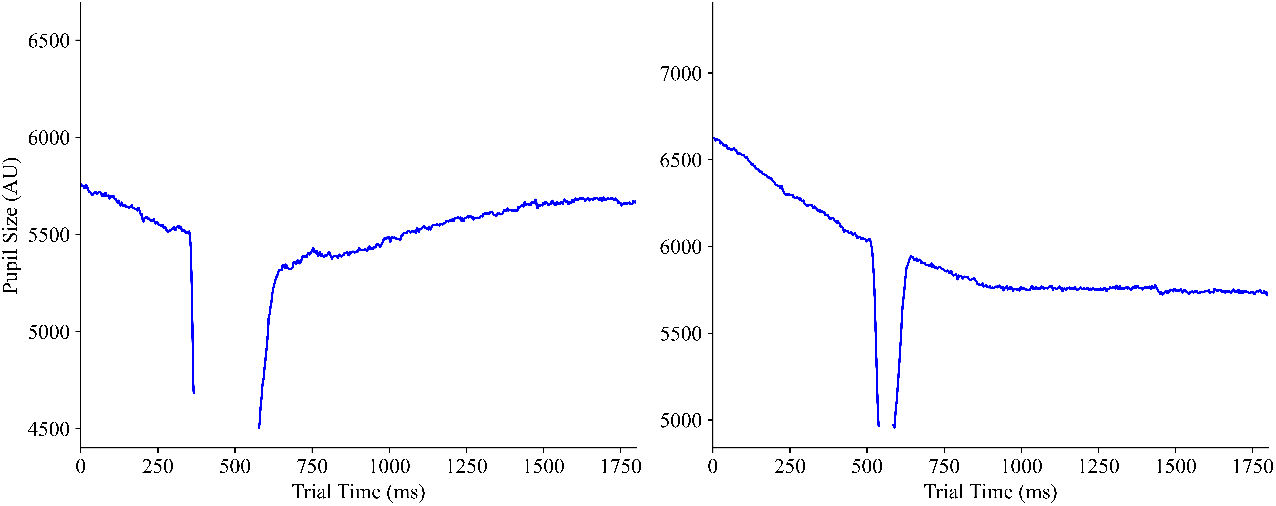
Visualization of two blink events leading to gaps in pupil size data. Regions corresponding to blink occurrences are empty, reflecting the absence of pupil size measurements.

**Fig. 2.**
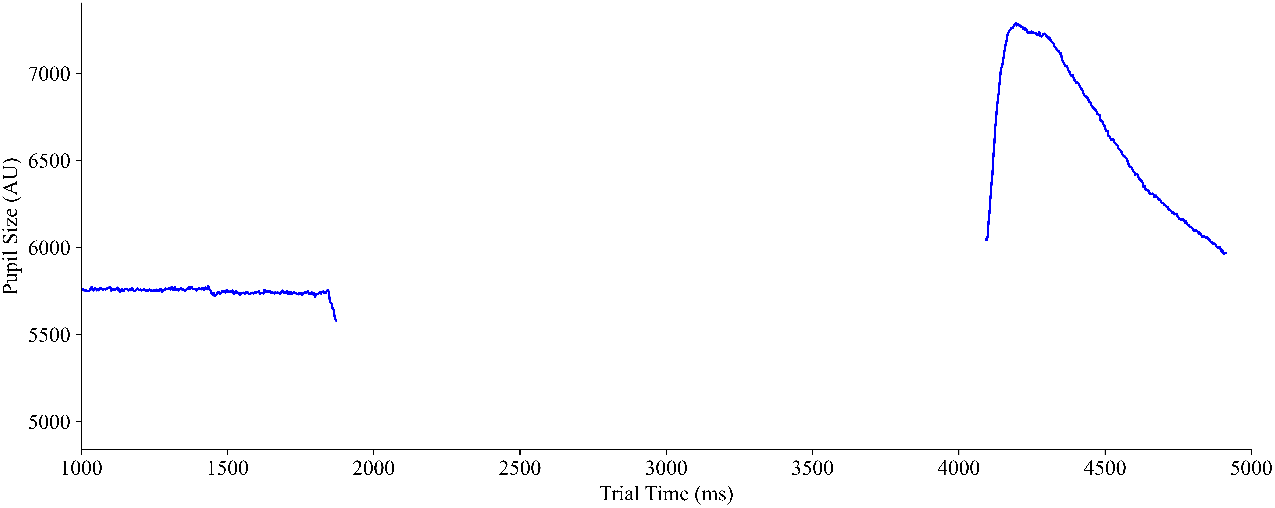
A long-duration blink where the eye tracker lost the pupil signal from approximately 1900 ms to 4100 ms. Note that this happened due to the participant’s looking at the keyboard to type responses to a task, “Reference: Author”

Blinks disrupt the recording process, causing missing or compromised data segments [22]. Such interruptions introduce noise, mask key behavioral patterns, and reduce analysis reliability by creating data gaps [16, 23] and artifacts [24], especially in studies that require continuous and accurate measurements of pupil size or fixation points.

Various methods have been developed to address blinks in eye-tracking data, each with varying degrees of sophistication. The most straightforward approach excludes affected data segments [25]. Although simple, this leads to the loss of valuable information and can introduce biases, mainly if blink events are not uniformly distributed [26–28]. More advanced methods include velocity-based algorithms [29], which infer blink onsets and offsets from pupil size velocity changes, and noise-based techniques [22, 30] that identify blink-related noise patterns. Pedrotti [31] introduced a data-driven method to correct blink-related artifacts within pupil diameter measurements. In particular, Hershman et al. [22] proposed an algorithm that uses device-specific noise characteristics to detect blink boundaries. Despite these advancements, many methods remain constrained by a lack of adaptability [22], with researchers often implementing ad-hoc pipelines. These range from ignoring missing data to artificial interpolation, which can degrade the accuracy of subsequent analyses [32]. Moreover, low-speed eye tracking protocols complicate accurate event detection, affecting higher-level analyses [31].

The limitations of existing blink detection methods highlight the need for more robust and generalizable approaches. Currently, many techniques are based on predefined thresholds [31] or device-specific noise profiles [22], compromising the effectiveness in diverse data sets. Subtle, short-duration, and overlapping blinks often result in inaccuracies in onset and offset detection [33, 34]. Additionally, edge cases remain problematic, for example, when recordings begin or end with missing data, making it difficult to refine the boundaries of blink detection [35]. These challenges underscore the importance of developing more flexible methods that detect blinks without relying on rigid device-specific assumptions.

To address these challenges, this study introduces a novel blink detection model that enhances reliability across diverse experimental conditions. The model incorporates dynamic thresholding (see Section A), Gaussian smoothing (see Secttion B), and a trend-based boundary refinement (see Section C). Unlike conventional approaches that rely on fixed thresholds or device-specific noise patterns, this model adapts dynamically to pupil size fluctuations, ensuring robust performance across different eye-tracking systems and experimental designs. By treating each blink as an independent event, it preserves temporal resolution, making it suitable for cognitive neuroscience, fatigue monitoring, and clinical pupillometry research. This provides researchers with a versatile and computationally efficient tool for handling blink-induced data disruptions, ultimately improving the integrity of eye-tracking analyses

## 2 Methods

The proposed model first applies a dynamic thresholding strategy to establish a robust criterion for identifying missing data segments. This approach derives a threshold as a fraction of the mean pupil size computed on nonzero samples. This threshold is mathematically defined as:

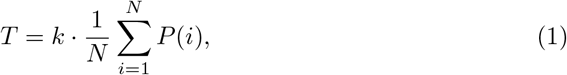

where *P* (*i*) denotes the pupil size at sample *i, N* is the number of valid (non-zero) samples, and *k* is a threshold ratio, typically ranging between approximately 0.2 and 0.5 [29, 36]. The threshold *T* is dynamically calibrated based on the mean pupil size of valid samples, ensuring robustness across datasets where pupil size reductions may indicate blinks even if the pupil does not reach zero. The threshold ratio *k* acts as a scaling factor, allowing detection sensitivity to be adjusted for different experimental conditions and participants. This approach prevents over-sensitivity in large pupils and missed blinks in smaller pupils. By defining the threshold *T* as a fraction of the mean pupil size, the model ensures that detection is not affected by absolute differences in pupil size across individuals or devices. (for more details, see Section D, Figure 3).

**Fig. 3.**
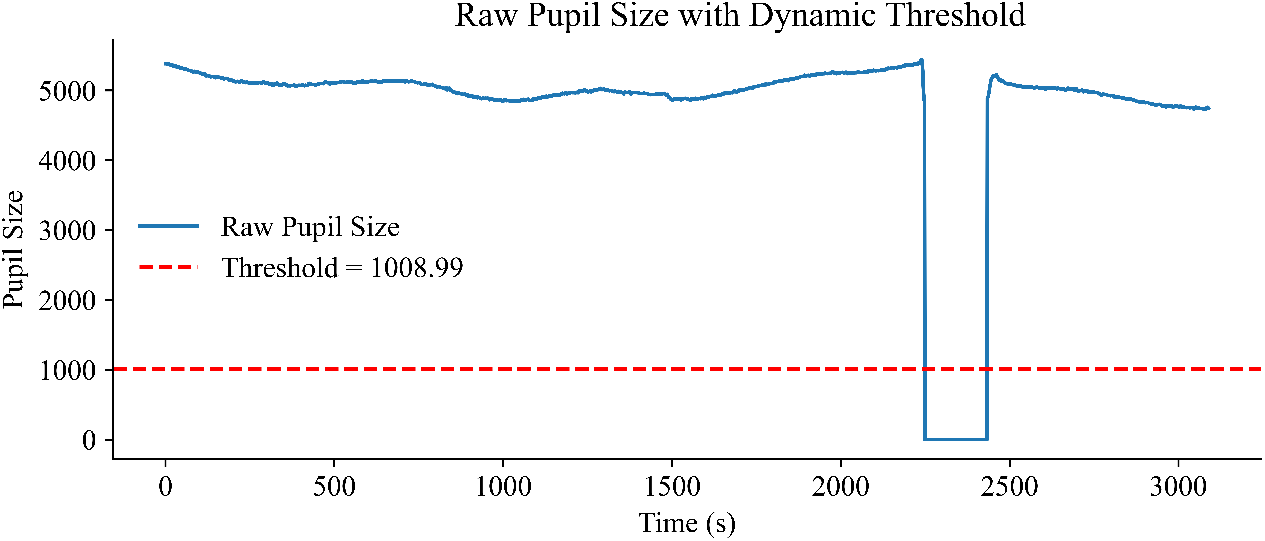
Dynamic thresholding establishes an adaptive threshold for detecting blink events based on pupil size distribution

With the threshold established, the next step constructs a binary indicator signal *M* (*t*) to mark the intervals of missing data; this method is presented by [22] and these intervals reserve the data points for further steps. Here, *M* (*t*) equals 1 if the sample is considered missing (i.e., below the threshold) and 0 otherwise:

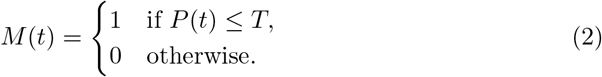

This binary representation delineates between intact and missing data segments, enabling systematic detection of blink events as shown in Figure 4.

**Fig. 4.**
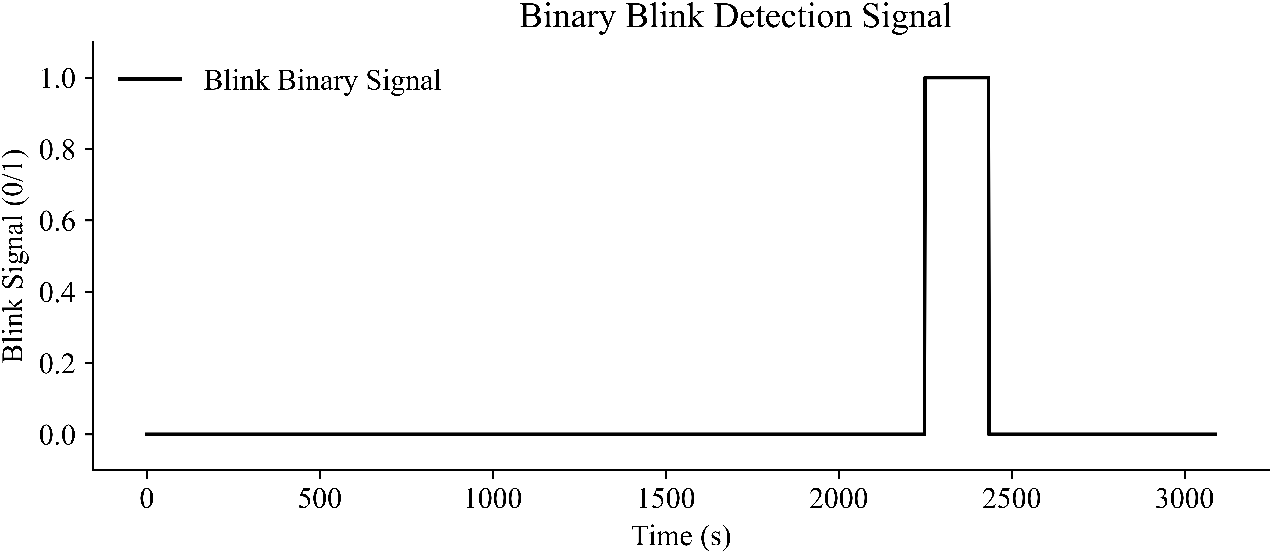
The binary indicator signal marks missing data points associated with blink events, distinguishing them from valid pupil size measurements.

Transitions in *M* (*t*) are analyzed using forward differencing to identify the onset and offset of blinks:

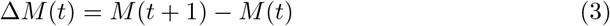

where a positive change (Δ*M* (*t*) = 1) corresponds to a blink onset, and a negative change (Δ*M* (*t*) = −1) marks a blink offset. This provides the initial delineation of blink intervals (Figure 5).

**Fig. 5.**
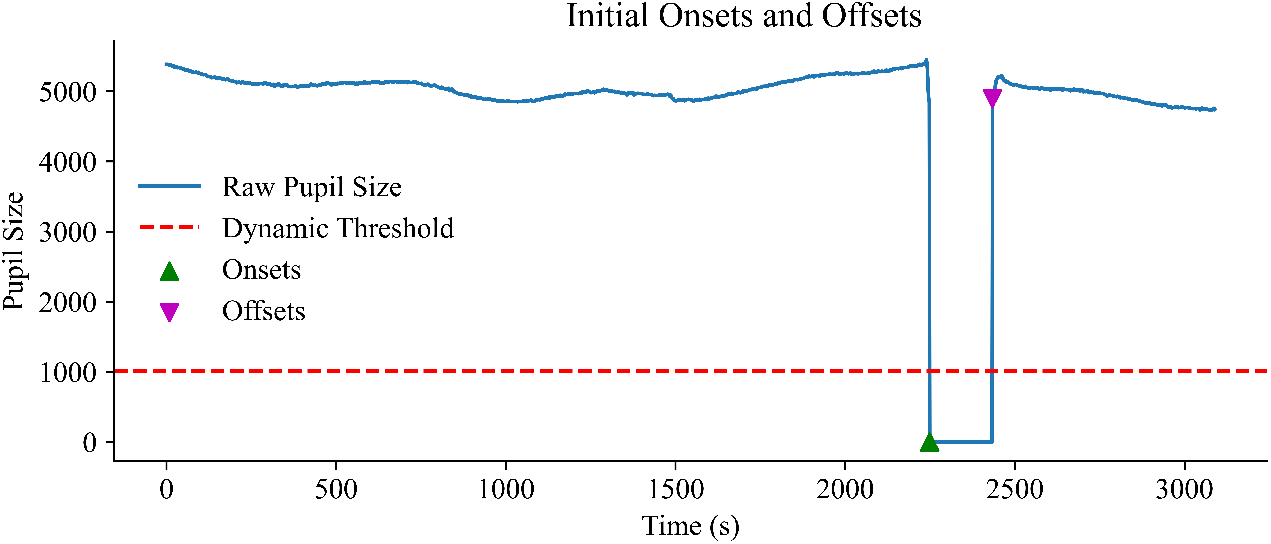
Initial detection of blink onsets and offsets. These preliminary values may be further refined in subsequent steps to improve accuracy

The raw pupil size signal undergoes Gaussian smoothing to refine these initial estimates further [36–39]. This step reduces noise while preserving critical transitions, improving the clarity of onset and offset boundaries. The smoothed signal *P*_smooth_(*t*) is obtained using convolution with a Gaussian kernel:

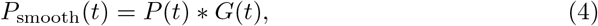

where *G*(*t*) represents the discrete Gaussian kernel. Filtering out high-frequency fluctuations facilitates more accurate boundary detection (Figure 6).

**Fig. 6.**
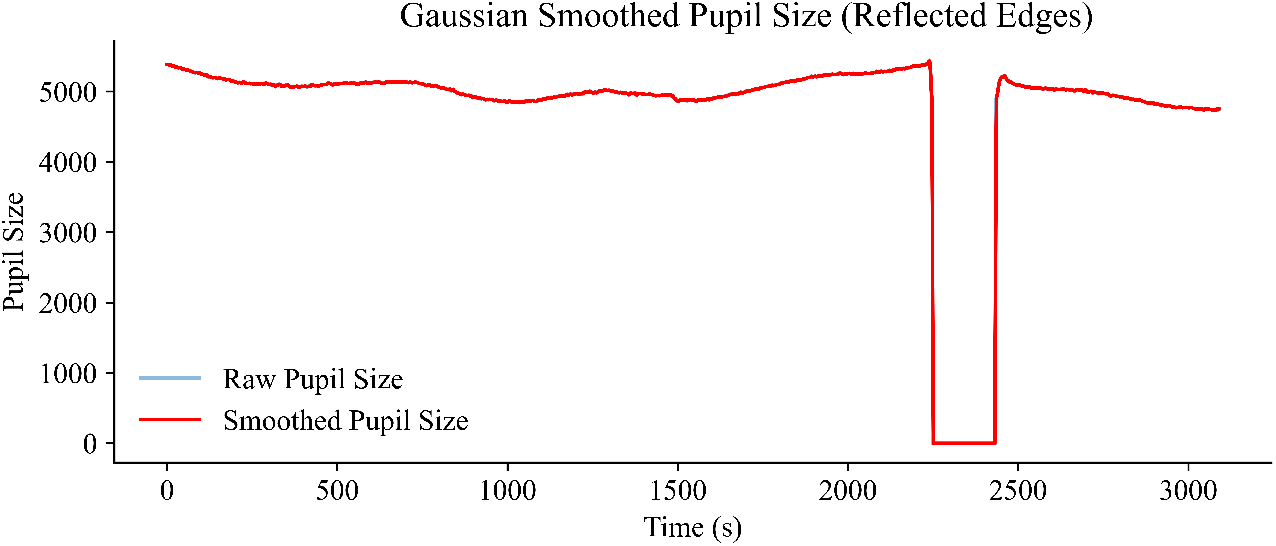
Gaussian smoothing reduces high-frequency noise while preserving essential signal characteristics, enhancing the accuracy of detected blink boundaries.

Next, the detected onsets and offsets are refined by tracking trends in the smoothed signal as shown in the Figure 7. This ensures that the final blink boundaries align closely with meaningful physiological changes rather than transient noise. The refined onset *t*_onset_ is determined by searching backward from the initially detected onset until the smoothed signal stabilizes at a low level:

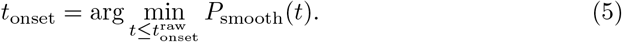

**Fig. 7.**
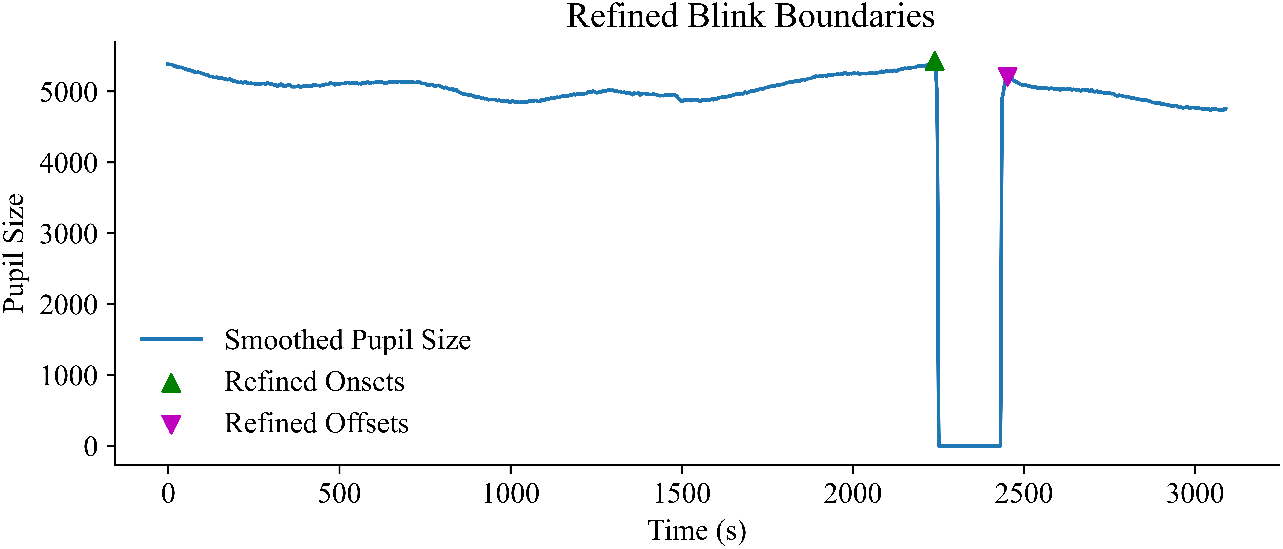
Refinement of blink boundaries using a trend tracking.

Similarly, the refined offset *t*_offset_ is found by moving forward from the raw offset:

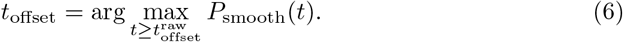

Unlike some alternative methods that merge closely spaced blinks into a single event, this model treats each blink as an independent event. This is particularly important for studies analyzing blink frequency, inter-blink intervals, or temporal dynamics, ensuring that every closure is counted as a discrete event. and finally as an optional illustration you can see the final detected area as blink in the Figure 8

**Fig. 8.**
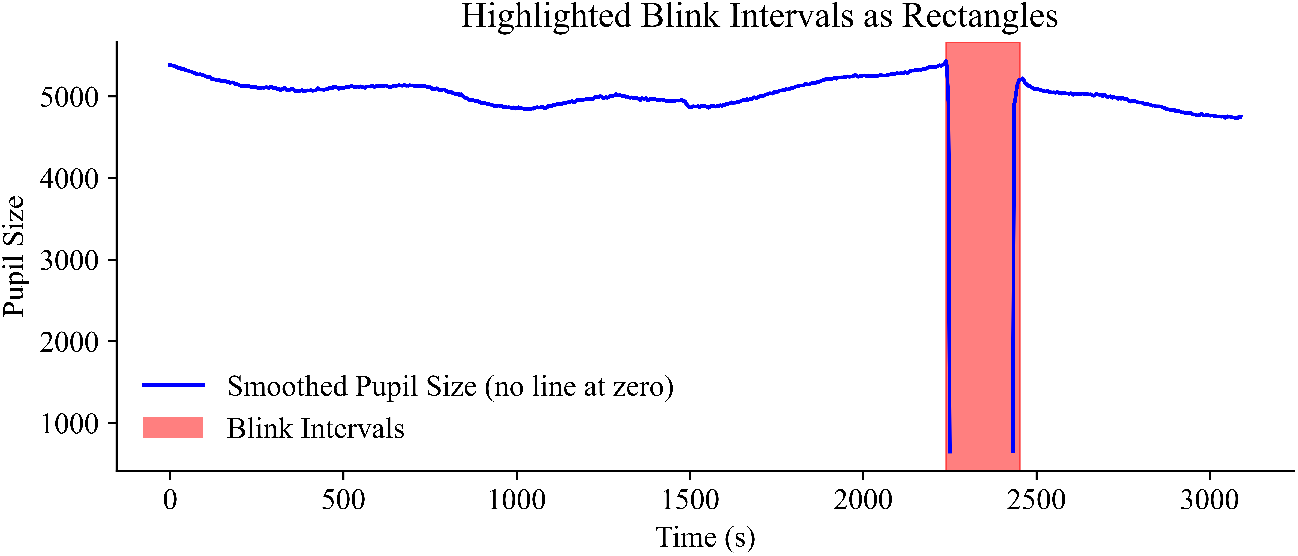
Final delineation of blink events. In this section, the detected blink area is shown in red color.

Figure 9 shows an example of the model successfully detecting and delineating blink onsets and offsets, resulting in a cleaner and more interpretable signal. In addition, Figure 10 provides a more challenging scenario, demonstrating the robustness of the model in handling extreme conditions. Even when faced with atypical blink patterns—ranging from extremely short to unusually long signal losses due to participant movement or gaze deviation—the model adaptively refines blink boundaries, maintaining reliable detection performance across a broad spectrum of recording conditions.

**Fig. 9.**
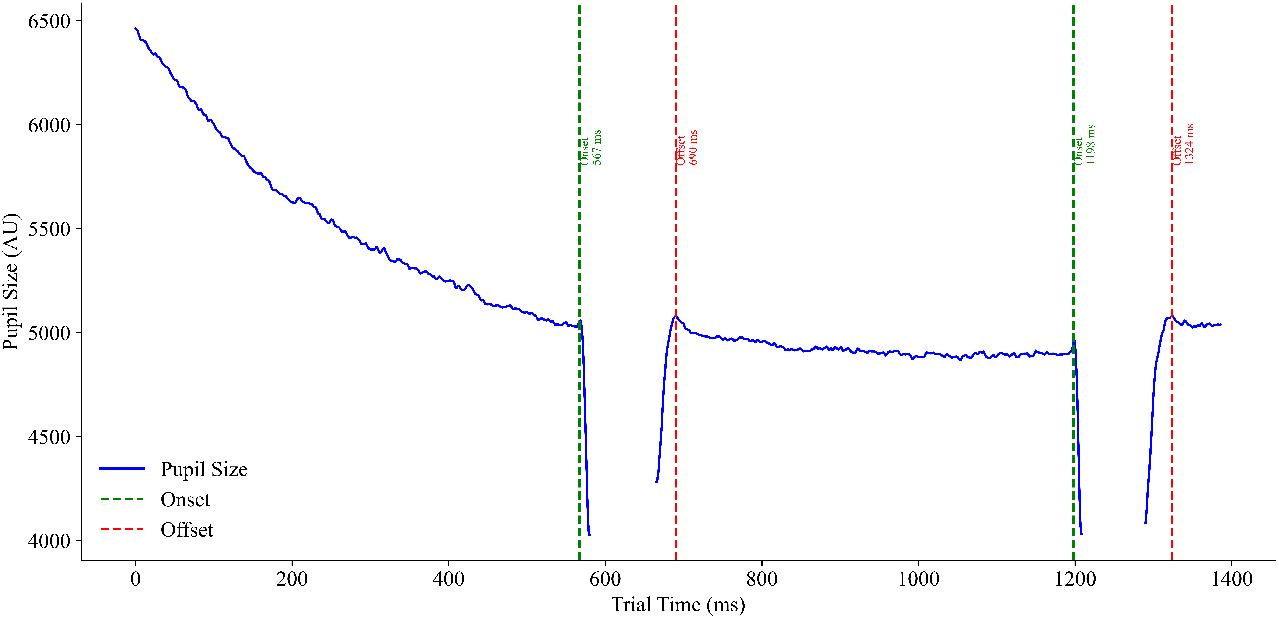
Demonstration of the model-based blink onset and offset detection. The plot illustrates how the model accurately identifies blink events within the pupil size signal, ensuring clear delineation of blink boundaries.

**Fig. 10.**
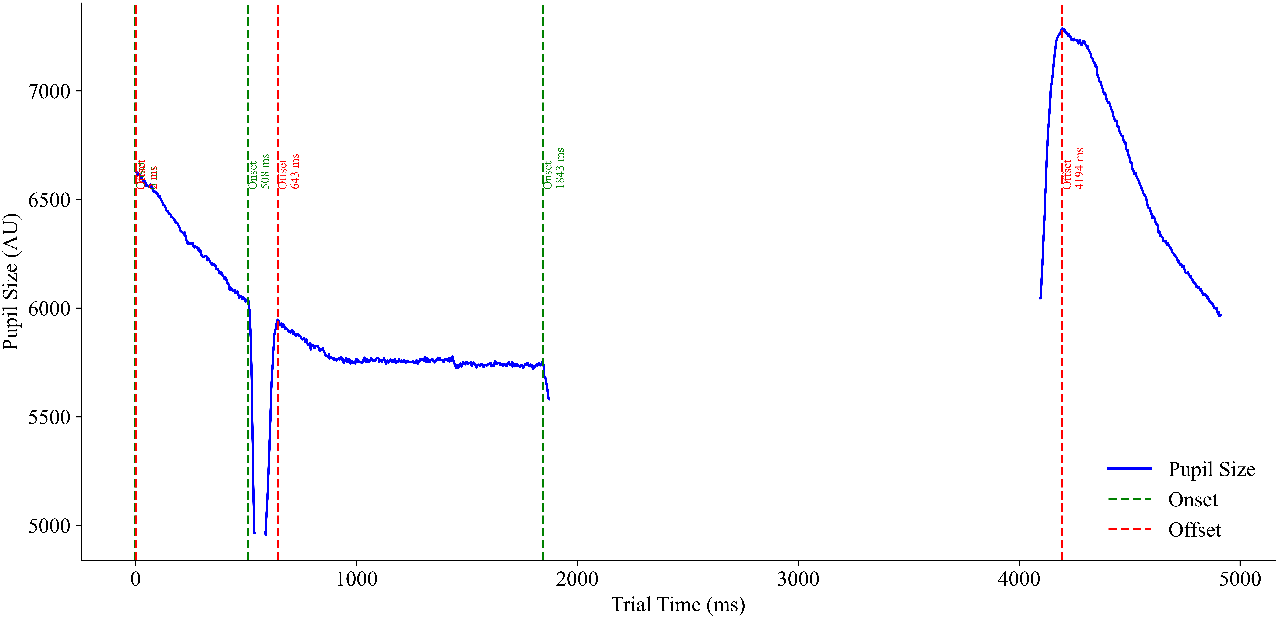
An extreme case involving multiple blink events. The model detects a very short blink (offset at 2 ms), a typical blink (508–643 ms), and a prolonged blink (1843–4194 ms) caused by participant movement or gaze redirection. This example highlights the model’s flexibility and robustness in adapting to diverse and challenging data conditions.

Finally, all steps are taken together; Figure 11 shows the plot of all steps done with the function.

**Fig. 11.**
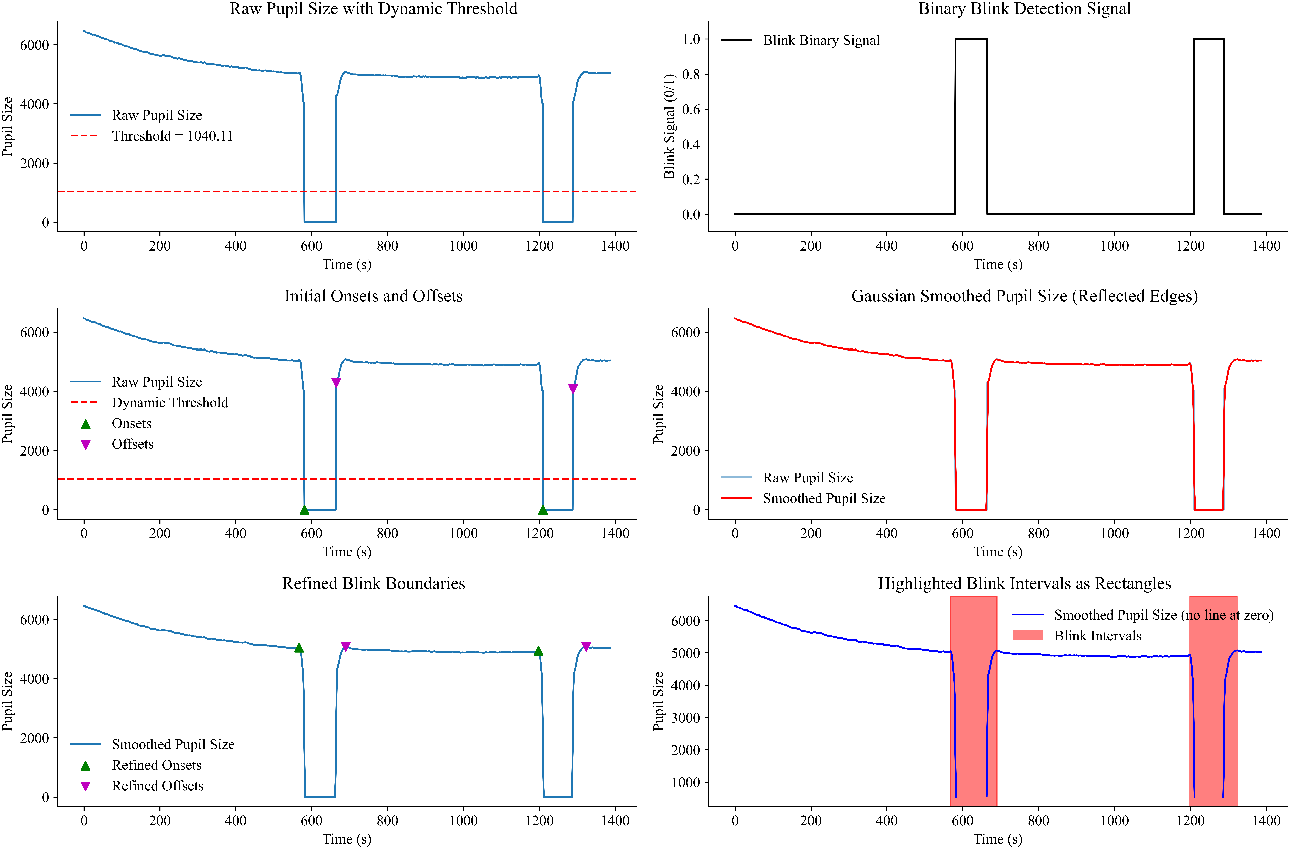
This figure illustrates the complete blink detection pipeline, presented in a sequence of steps arranged in a 3*×* 2 subplot format. In Step 1, a dynamic threshold is computed based on the non-zero pupil size values, providing a baseline against which potential blinks are assessed. In Step 2, a binary signal highlights intervals where pupil size falls below this threshold, indicating potential blink periods. In Step 3, initial blink onsets and offsets are identified as transitions in the binary signal, providing a preliminary segmentation of blink events. In Step 4, Gaussian smoothing with reflective padding is applied to reduce noise and minimize edge artifacts in the pupil size data. Building on this, Step 5 refines the blink boundaries by shifting onsets and offsets to ensure they align with the smooth signal’s physiologically plausible transitions. Finally, in Step 6, zeros are replaced with NaNs to prevent line continuation through blink periods, and the detected blink intervals are visually emphasized as shaded red rectangles, clearly delineating the final blink events.

### Advantages of the Proposed Model

The proposed blink detection model introduces several key improvements over traditional methodologies, enhancing its applicability and reliability in pupillometry research. A central innovation of the model is its dynamic thresholding mechanism, which determines missing data based on a fraction of the mean pupil size of valid samples. Unlike fixed-threshold approaches that assume blinks correspond strictly to zero pupil size, this method adapts to interindividual differences and baseline fluctuations by setting a relative threshold. This adaptability allows for greater robustness across datasets with varying noise levels, ensuring that both full and partial blinks—where pupil size significantly decreases but does not reach zero—can be detected.

A second major improvement is the incorporation of Gaussian smoothing, which effectively reduces noise while preserving key signal features [40]. Unlike simple averaging filters, Gaussian smoothing assigns greater weight to central data points within the window, minimizing artifacts and preventing distortions introduced by abrupt signal changes. This approach is particularly beneficial in datasets with high-frequency fluctuations or measurement noise [21, 41], enabling more precise delineation of blink onsets and offsets.

Another distinguishing feature of the model is its refinement strategy for blink boundaries. Instead of relying solely on abrupt pupil size changes, the model iteratively refines blink onsets and offsets by tracking trends in the smoothed pupil signal. This ensures that detected blink intervals align closely with physiological eyelid movements, reducing false positives while maintaining sensitivity to short-duration or subtle blinks that conventional methods may overlook.

Unlike methods that merge consecutive blinks based on a fixed time threshold, our approach preserves inter-blink intervals, allowing for a more accurate estimation of blink frequency and temporal dynamics. This is particularly important in fatigue detection studies, where blink rate variability provides key insights into cognitive state fluctuations. For instance, some models, such as that of [22], concatenate closely spaced blinks into a single event, potentially obscuring rapid fluctuations in blink activity. In contrast, our model treats each blink individually, ensuring that even closely spaced blinks are separately identified. This is critical for research contexts where precise temporal resolution is required, such as in studies with high-speed visual stimuli (e.g., presented in 10 ms intervals). In such cases, blink merging may artificially reduce the number of detected events, leading to potential information loss in time-sensitive experiments. By maintaining blink independence, our model ensures that each eyelid closure is recorded as a distinct event, thereby preserving the integrity of temporal analyses without artificially modifying blink counts.

Additionally, the model is designed to handle challenging cases, such as blinks occurring at the start or end of the recording without misclassification. Instead of enforcing artificial constraints on boundary cases, the method detects blinks based solely on the pupil size signal, ensuring consistency across different experimental conditions.

Finally, the model is computationally efficient, leveraging lightweight operations such as forward differencing and Gaussian smoothing to achieve accurate blink detection with minimal processing overhead. This efficiency makes it suitable for high-throughput applications and real-time or near-real-time pupillometry studies. By balancing detection accuracy with computational efficiency, the proposed method provides a scalable solution for both laboratory research and applied settings where blink monitoring is critical.

## 3 Discussion

This study introduces a novel blink detection model that overcomes the limitations of existing pupillometry approaches. The model significantly improves robustness, precision, and adaptability by integrating dynamic thresholding, Gaussian smoothing, and a trend-based onset/offset refinement. Such improvements render it a powerful analytical tool, particularly in real-world contexts where noise, pupil size variability, and complex edge cases impede the accuracy of conventional methods.

A key contribution of this model lies in its ability to generalize across heterogeneous datasets. Rather than relying on fixed thresholds or device-specific noise profiles, the dynamic thresholding mechanism of the model adaptively aligns with the intrinsic characteristics of the data, ensuring reliable detection of both subtle and pronounced blink events. This adaptability, combined with the model’s a trend-based refinement, reduces false positives and enhances the physiological validity of detected blinks.

The incorporation of Gaussian smoothing constitutes a critical preprocessing step, effectively minimizing noise while preserving the underlying structure of the pupil size signal. This approach sharpens the distinction between authentic blinks and spurious artifacts in datasets characterized by high-frequency fluctuations. By leveraging adjustable parameters such as threshold ratio and smoothing window size, the model can be fine-tuned to meet various experimental demands, extending its utility to cognitive neuroscience, fatigue assessment, and clinical diagnostics.

Unlike some alternative approaches that merge closely spaced blinks into a single event, this model treats each detected blink individually. This is particularly important for research contexts where blink frequency, inter-blink intervals, or temporal dynamics are of interest. By maintaining blink independence, the model ensures that each closure is recorded as a discrete event, preserving the integrity of temporal analyses without artificial modifications to blink counts.

Additionally, the model is designed to handle challenging cases such as blinks occurring at the start or end of the recording without misclassification. Rather than enforcing artificial constraints on boundary cases, the method detects blinks based solely on the pupil size signal, ensuring consistency across different experimental conditions.

Despite these advances, certain limitations remain. Although beneficial for flexibility, the dependence of the model on user-defined parameters, such as the threshold ratio and smoothing window size, necessitates careful selection based on the specific dataset characteristics. Future work could explore adaptive thresholding techniques that automatically adjust based on signal properties, further enhancing robustness across diverse experimental contexts. Additionally, while Gaussian smoothing has demonstrated strong noise-reduction capabilities, investigating alternative smoothing techniques may further optimize performance in datasets exhibiting extreme variability.

While this method improves robustness over static thresholding approaches, it still requires manual selection of parameters such as the threshold ratio *k* and smoothing window size. Future work could explore data-driven adaptive parameter selection methods to further enhance robustness across diverse datasets. Additionally, while Gaussian smoothing improves onset/offset accuracy, alternative signal processing techniques, such as wavelet-based denoising or machine-learning-based segmentation, may provide further refinements in detecting complex blink dynamics.

In conclusion, this work presents a robust and adaptable model that transcends the limitations of traditional blink detection methods. The model provides a reliable and efficient solution for analyzing eye-tracking data across numerous research domains by incorporating dynamic thresholding, Gaussian smoothing, and trend-based refinements. Its resilience to real-world challenges and computational efficiency make it valuable for diverse scientific investigations. As parameter optimization strategies evolve and further validations against diverse datasets unfold, the utility of this model will continue to expand, shaping the future of pupillometry and related fields.

## Supplementary information

The Python code used for data analysis and visualization, including the implementation of the current model and associated plotting functions, is freely available at (https://osf.io/bgqwv) as a single function code, which implies the model is available at (https://osf.io/689xd). In addition, the code for the plotting steps is available at (https://osf.io/gy9b7).

## Acknowledgements

The author would like to express profound gratitude to Professor Luca Battaglini from the University of Padua; Yash Vyas, from Department of Industrial Engineering, University of Padua; Professor Jeremy Wolfe from the Visual Attention Lab at Harvard University; Professor Michele Rucci from the University of Rochester; Professor Eghbal Hosseini from the Technical University of Denmark; and Oryah Lancry from Visual Attention Lab; In addition, heartfelt appreciation is extended to Harvard Visual Attention Lab colleagues for their invaluable contributions to this work.

## Data Availability

Code and Data used for this work are avaibale at the OSF repository at https://osf.io/xpdce/

## Appendix A Why Dynamic Threshold

Dynamic thresholding, or calibrating the detection cutoff according to the statistical properties of the dataset, offers a more resilient alternative to fixed thresholding in pupillometry because it accounts for natural fluctuations in baseline pupil size and measurement conditions [42, 43]. Instead of applying an absolute threshold that may fail when faced with inter-individual differences [44, 45] or alterations in lighting and equipment, dynamic thresholding continuously adapts to the overall pupil size distribution of each data set by referencing the mean or median of valid samples [42, 43, 46]. This approach ensures that even moderate drops in pupil size, which may indicate partial eyelid closure or incomplete blinks, are adequately detected without overestimating noise-related artifacts. Researchers such as Hershman et al. [22] emphasize the need to consider data- and device-specific noise characteristics and caution that static thresholds often overlook subtle details in real-world settings. Kret and Sjak-Shie [36] highlight the importance of normalization and adaptive parameterization when identifying blinks, noting that extreme uniformity in thresholding criteria can lead to inconsistencies between participants and instruments. Sirois and Brisson [29] also underscore the benefits of adaptive approaches that scale with the data rather than imposing rigid cut-offs because pupil size measurements inherently vary across time [20], tasks [47], and individuals [48, 49]. By tuning the threshold to the central tendencies of each data set, the method remains sensitive to blink-related fluctuations while reducing false detections arising from measurement artifacts or transient changes in gaze.

## Appendix B Gaussian smoothing

Gaussian smoothing mitigates high-frequency fluctuations and measurement artifacts [50, 51] in the pupil size signal by applying a convolution filter with a Gaussian kernel. This filter weights data points within a local window, giving more influence to central values, thereby producing a smooth curve with reduced abrupt jumps [52–55]. This process helps preserve essential low-frequency features likely to reflect genuine physiological changes, such as the transitions between blink and non-blink intervals, while attenuating random noise arising from device precision limits, participant micro-movements, or environmental factors. By smoothing out rapid spikes or dips in pupil size that do not represent actual events, the technique improves the reliability of onset and offset detection in blink analysis, allowing the subsequent detection algorithm to base its decisions on more stable patterns in the data. The preserved structural integrity of the underlying signal is fundamental in pupillometry, where slight variations can indicate nuanced cognitive or physiological states and where the risk of mistaking these variations for actual blinks is heightened if the raw signal contains too much noise.

## Appendix C Flexible boundary refinement

The model refines detected blink onsets and offsets by leveraging a trends in the smoothed pupil signal. Instead of relying on strict threshold-based cutoffs, the approach shifts blink boundaries to ensure they align with stable transitions in the smoothed data. This refinement step helps mitigate noise-driven errors and prevents premature or delayed marking of blink events.

For an initially detected onset 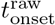, the refined onset *t*_onset_ is determined by scanning backward in time until a stabilization point is reached, ensuring the onset aligns with the minimum pupil size before the blink event:

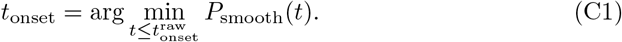

Similarly, the refined offset *t*_offset_ is determined by searching forward in the smoothed signal until a stabilization point is reached, ensuring it aligns with the maximum pupil size after the blink event:

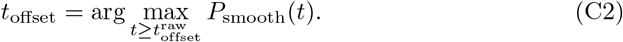

By dynamically adjusting blink boundaries based on the smoothed signal, this approach reduces the risk of misclassification caused by transient noise or abrupt signal fluctuations. Unlike static methods, which may set arbitrary fixed thresholds for refinement, this adaptive technique allows for better physiological alignment with real blink events.

## Appendix D Selecting threshold ratio

### Logic Behind Using a Fraction *k* of the Mean Pupil Size

The decision to define a blink detection threshold *T* as a fraction *k* of the mean pupil size is based on the need for a relative data-driven criterion that adapts to varying conditions and individual differences in the size of the baseline pupil. Rather than relying on a fixed absolute value, this approach ensures that the threshold scales with the intrinsic characteristics of the dataset. The following are the key points behind this logic.

Different participants and experimental conditions can influence the baseline pupil sizes [47–49]. For instance, one participant may have a naturally larger pupil or lighting conditions may shift the average pupil diameter. Using a single absolute threshold for all datasets and conditions may lead to misclassification, as what constitutes a blink-level drop in one dataset might not apply to another.

By setting *T* as a fraction of the mean pupil size, the threshold dynamically scales with individual pupil size variations. This ensures that larger baseline pupils require a proportionally higher drop for blink detection, while smaller baseline pupils require a lower drop. Selecting *k* within this range is informed by practical considerations [29, 36]. Typically, blink events manifest as significant reductions in pupil size [56], often close to zero. However, measurement noise or partial closure of the eyelid may produce values that are not strictly zero, but still represent a pronounced drop [57, 58]. In which A lower *k* (e.g., 0.2) is more lenient, capturing subtle drops and ensuring greater sensitivity to slight decreases in pupil size, and a higher *k* (e.g., 0.5) is stricter, requiring more substantial decreases in pupil size for detection, thus reducing false positives. Adjusting *k* provides a means to fine-tune the sensitivity and specificity of blink detection. A threshold ratio within 0.2–0.5 often serves as a balanced starting point, offering broad applicability while allowing for experiment-specific refinements [59].

While there is no universally fixed threshold ratio, using a fraction of the mean pupil size has emerged from empirical observations and discussions in pupillometry. Researchers have consistently observed substantial variability between participants, conditions, and measurement devices. Consequently, they have advocated for data-driven techniques that derive thresholds from the dataset’s distribution of pupil sizes rather than imposing absolute criteria.

In other words, this range is neither too restrictive nor too permissive, accommodating subtle blink-related drops without over-flagging minor fluctuations. Such empirical insights have guided practitioners toward using a relative threshold that harnesses the inherent characteristics of the data, making the blink detection process both more robust and more tailored to the specifics of each dataset.

